# High ambient temperature facilitates the acquisition of 3,4-methylenedioxymethamphetamine (MDMA) self-administration

**DOI:** 10.1101/123828

**Authors:** Shawn. M. Aarde, Pai-Kai Huang, Michael A. Taffe

## Abstract

**Rationale:** MDMA alters body temperature in rats with a direction that depends on the ambient temperature (T_A_). The thermoregulatory effects of MDMA and T_A_ may affect intravenous self-administration (IVSA) of MDMA but limited prior reports conflict.

**Objective:** To determine how body temperature responses under high and low T_A_ influence MDMA IVSA.

**Methods:** Male Sprague-Dawley rats were trained to IVSA MDMA (1.0 mg/kg/infusion; 2-hr sessions; FR5 schedule of reinforcement) under T_A_ 20°C or 30°C. Radiotelemetry transmitters recorded body temperature and activity during IVSA.

**Results:** MDMA intake increased under both T_A_ during acquisition, but to a greater extent in the 30°C group. The magnitude of hypothermia was initially equivalent between groups but diminished over training in the 30°C group. Within-session activity was initially lower in the 30° C group, but by the end of acquisition and maintenance, activity was similar for both groups. When T_A_ conditions were swapped, the hot-trained group increased MDMA IVSA under 20 °C T_A_ and a modest decrease in drug intake was observed in the cold-trained group under 30 °C T_A_. Subsequent non-contingent MDMA (1.0-5.0 mg/kg, i.v.) found that rats with higher MDMA IVSA rates showed blunted hypothermia compared with rats with lower IVSA levels; however, within-session activity did not differ by group. High T_A_ increased intracranial self-stimulation thresholds in a different group of rats and MDMA reduced thresholds below baseline at low, but not high, T_A_.

**Conclusions:** High T_A_ appears to enhance acquisition of MDMA IVSA through an aversive effect and not via thermoregulatory motivation.

## 1. Introduction

Recreational use of (±)3,4-methylenedioxymethamphetamine (MDMA; “Ecstasy”) became increasingly popular in recent decades [1-3]. In the USA, annual prevalence rates have been around 5% of respondents for young adults in the past decade [4-6]; *lifetime* prevalence for Ecstasy of 11-12% are reported in recent years. Annual prevalence of Ecstasy exposure is at least 3 fold higher than prevalence for heroin, crack (smokable cocaine), ice (smokable methamphetamine) or PCP. Thus, overall lifetime rates of exposure to Ecstasy are substantial and will continue to be so for some time, until these cohorts expire. It is further concerning that multiple Phase I clinical trials are underway to establish MDMA as an adjunctive treatment for psychotherapy [7-12] because surveys of attitudes toward drug risk show that diverted pharmaceutical preparations (e.g., amphetamines) have a perception of safety which coincides with higher incident rates [4, 5, 13]. Increased population exposure is problematic because significant proportions of heavy Ecstasy users meet criteria for dependence at some point in their use history [3, 14-16]. There are also case reports of Ecstasy/MDMA use patterns that are daily or at least several times per week [17-19].

Laboratory studies of the abuse liability of MDMA have been curiously sporadic in comparison with many other drugs of abuse. MDMA will substitute for cocaine in baboons and rhesus monkeys trained for intravenous self-administration [20-24] and it generates consistent, but low levels of self-administration in rats [25-27]; one laboratory reports intakes at least several fold higher [28-30]. The Schenk lab has reported that only about 60% of rats reach acquisition criteria although additional animals may acquire after 21+ days of access [31-33]. There are also hints of increased MDMA intake with many sessions of access [25, 34, 35] or with longer daily access sessions [36].

A prior observation reported up to twice as many infusions of MDMA were self-administered in rats when ambient temperature was increased to 30°C in single day challenges after initial training under normative (21°C) ambient temperature [37]. A similar phenomenon may also occur in nonhuman primate models [38]. This may be of critical translational importance since many human Ecstasy users ingest the drug in the context of a crowded dance club environment. This finding may also explain inconsistency in prior rodent investigations of MDMA self-administration. Rats’ body temperature is decreased by MDMA at low ambient temperature and increased at high ambient temperature [39-41] which may be behaviorally motivational [42-44]. The physiologically thermoneutral range for rats is around 30°C, but behavioral preference in a thermocline may be some 6-8°C lower [45], although also see [46] for behavioral preference for 27-29°C. This raises the possibility that behavioral motivation may be altered by temperature decreases caused by self-administered MDMA at normal laboratory temperatures, typically below ∼25°C. Consequently the results of Cornish and colleagues (2003) could potentially be explained by an asymmetry in the aversion to body temperature responses to MDMA under different ambient temperatures.

This study was designed to determine if intravenous MDMA self-administration is increased in rats by training in a high ambient temperature and if so, if this is related to a change in the thermoregulatory response to intravenously self-administered MDMA. The threshold for thermoregulatory effect of acute MDMA appears to be 5 mg/kg [40] and it is unknown if various intravenous self-administration patterns [0.5-1.0 mg/kg/inf is typical, resulting, e.g., in 4-7 mg/kg over 2 hrs [27], 30 mg/kg over 6 hrs [32] and 30 mg/kg over 2 hrs [33]] alter body temperature. A minimally invasive implanted radiotelemetry system, previously shown sensitive to the temperature disrupting effects of non-contingent administration of Δ9-tetrahydrocannabinol, mephedrone (4-methylmethcathinone), 3,4-methylenedioxypyrovalerone (MDPV), α-pyrrolidinopentiophenone (α-PVP) and MDMA [47-50] as well as the intravenous self-administration of 4-methylmethcathinone [51] in rats, and similar to that found sensitive to effects of MDMA, methamphetamine and THC in monkeys [52-55], was used to minimize behavioral disruption during the self-administration sessions. Finally, the effect of high and low ambient temperature on the effects of MDMA on intracranial self-stimulation reward was determined to test the hypothesis that high ambient interferes with the ability of MDMA to reduce brain reward thresholds.

## 2. METHODS

### 2.1 Animals

Male Sprague-Dawley (Harlan, Livermore, CA; Experiment 1: N = 24) and Wistar rats (Charles River; Experiment 2, N = 10); were housed in humidity and temperature-controlled (23±1 °C) vivaria on 12:12 hour light:dark cycles. Animals entered the laboratory at 10-13 weeks of age and weighed 350-400 grams at the start of the study. Animals had *ad libitum* access to food and water, except for during pellet training and drug self-administration sessions, (see below). Procedures were conducted under protocols approved by the Institutional Care and Use Committees of The Scripps Research Institute and consistent with the NIH Guide for the Care and Use of Laboratory Animals.

### 2.2 Drugs

Racemic MDMA (3,4-methylenedioxymethamphetamine HCl; provided by U.S. National Institute on Drug Abuse) was dissolved in physiological saline to a concentration of 1.0 or 0.5 mg/kg/inf per 0.1 ml of solution.

### 2.3 Surgeries

#### 2.3.1 Intravenous Catheterization

Rats were anesthetized with an isoflurane/oxygen vapor mixture (isoflurane 5% induction, 1-3% maintenance) and prepared with chronic intravenous catheters as previously described [56-58]. Briefly, the catheters consisted of a 14-cm length of polyurethane based tubing (Micro-Renathane®, Braintree Scientific, Inc, Braintree MA, USA) fitted to a guide cannula (Plastics One, Roanoke, VA) curved at an angle and encased in dental cement anchored to an ∼3 cm circle of durable mesh. Catheter tubing was passed subcutaneously from the animal's back to the right jugular vein. Catheter tubing was inserted into the vein and tied gently with suture thread. A liquid tissue adhesive was used to close the incisions (3M™ Vetbond™ Tissue Adhesive; 1469SB).

A minimum of 7 days was allowed for surgical recovery. For the first three days of the recovery period, an antibiotic (cefazolin; 0.4 g/mL, 2.0 mL/kg, s.c.) and an analgesic (flunixin; 2.5 mg/mL, 2.0 mL/kg, s.c.) were administered daily. During testing and training the catheters were flushed with heparinized saline before sessions and heparinized saline containing cefazolin (100 mg/mL) after sessions.

Catheter patency was assessed nearly once a week after the last session of the week via administration through the catheter of ∼0.2 ml (10 mg/ml) of the ultra-short-acting barbiturate anesthetic Brevital sodium (1% methohexital sodium; Eli Lilly, Indianapolis, IN). Animals with patent catheters exhibit prominent signs of anesthesia (pronounced loss of muscle tone) within 3 sec after infusion. Animals that failed to display these signs were considered to have faulty catheters and were discontinued from the study.

#### 2.3.2 Radiotelemetry Probe Implantation

Sterile radiotelemetry transmitters (Data Sciences International; CTA-F40 or TA-F40) were implanted in the abdominal cavity thru an incision along the abdominal midline posterior to the xyphoid space [48]. Absorbable sutures were used to close the abdominal muscle incision and the skin incision was closed with tissue adhesive. Post-operative care and recovery time was the same as that for i.v. catheterization. Body temperature and activity were measured as previously described [47-49]. In brief, the temperature was sampled every five minutes and the activity reflects a rate (of arbitrary counts of movement per five minutes) of the transmitter across the receiver plate that is placed under the cage.

#### 2.3.3 Intracranial Self-stimulation Electrode Implantation

Rats were anesthetized with an isoflurane/oxygen vapor mixture (isoflurane 5% induction, 1-3% maintenance) and prepared with stimulation electrodes as described in (Markou and Koob 1992). A small incision (approximately 1.5-4 cm) was made through the skin, the muscle carefully pushed aside using a blunt instrument, and the skull cleaned with sterile swabs or gauze. A stainless steel bipolar electrode (0.25 mm) was aimed at the medial forebrain bundle and implanted stereotaxically (in mm; +5 incisor bar, -0.5 AP from Bregma; ± 1.7 ML, -9.5 DV from skull at Bregma). The electrode was anchored to the skull with four to six stainless-steel screws and dental cement. The incision was closed using veterinary adhesive and/or suture. Post-operative care and recovery time was the same as that for i.v. catheterization.

### 2.4 Experiment 1: Self-administration

#### 2.4.1 Self-administration General Procedure

Animals underwent the following four sequential phases (detailed below): Lever-press training, MDMA intravenous self-administration (IVSA) *baseline* (3 session, all under 25 °C T_A_), IVSA *acquisition* under high vs. low T_A_ conditions, IVSA *maintenance*, IVSA T_A_ *swap* then non-contingent MDMA administration (NCA). For each type of session, subjects were transported to an experimental room (illuminated by red light) and placed into operant boxes (Med Associates, St. Albans, VT) located inside sound-attenuating chambers. For IVSA and NCA, catheter fittings on the animals' backs were connected to polyethylene tubing contained inside a protective spring suspended into the operant chamber from a liquid swivel attached to a balance arm. MDMA doses were delivered via syringe pump. Major adjustments for weight differences were done with concentration changes and minor adjustments were made to infusion duration, keeping within 3.5-5 seconds per infusion. Each operant session started with the extension of two retractable levers into the chamber. Following each completion of the response requirement (response ratio), a white stimulus light (located above the reinforced lever) signaled delivery of the reinforcer and remained on during a 20-sec post-infusion timeout, during which responses were recorded but had no scheduled consequences. Sessions began in the second half of the vivarium dark cycle, save for a total of 14 individual sessions (balanced across treatment group) that were begun within the first hour of the light cycle due to unavoidable experimental exigencies.

#### 2.4.2 Lever-Press Training

Prior to catheterization surgery, rats were food-restricted (∼15 g chow/rat/day) and trained to press the reinforced lever (always left lever) for delivery of 45-mg food pellets (TestDiet, Richmond, IN; Cat.# 1811156) under a fixed-ratio, one-press-per-reinforcer schedule of reinforcement (FR1). Once advancement criterion was achieved (50 reinforcers earned), the ratio requirement was increased to FR2 then to FR5 (same advancement criteria). On average, this training required 4±1 days. Rats were returned to ad libitum feeding immediately after completion of training and 1-3 days before surgery.

#### 2.4.3 Drug Self-administration – Baseline and Acquisition

After recovery from surgery, animals were again food restricted to ∼17 g chow/rat/day provided in the home cage for an unrestricted interval of time. Subsequently, rats began a sequence two-hour self-administration sessions (run ∼5 days per week) in which drug infusions (1.0 mg/kg/inf; ∼0.1 ml/inf; ∼4 sec/inf) were made available on an FR5 schedule of reinforcement. This approach was taken so that learning of the operant response (selective lever pressing to obtain a reward) was not confounded with the reinforcer efficacy of MDMA. The training dose was selected based on the protocol of the lab that has produced the most MDMA IVSA to date [59]. Animals remained in the operant chamber for an additional hour after the end of the SA session (levers retracted) to fully characterize the body temperature response. For the 1^st^ three sessions, all rats were tested at an T_A_ of 25 °C and under food restriction; rats were returned to *ad libitum* feeding immediately after the 3^rd^ session to eliminate established effects of food restriction on psychostimulant drug reward [60-62] from the design. Given the variability in prior studies of MDMA self-administration it was deemed essential to balance the groups rather than rely on random assignment. For the next 14 sessions, rats were split into two groups (balanced for infusions over the 1^st^ three sessions); one group was trained in sessions under an T_A_ of 30 °C (“Hot-trained” group) and the other group trained under an T_A_ of 20 °C (“Cold-trained” group). Thirty minutes after the start of a session, a single priming infusion was given to any animals that had yet to self-administer an infusion in that session. A saline-substitution session immediately followed this Acquisition phase (see Saline Substitutions below). One rat that was assigned to train under 30 °C T_A_ was excluded due to catheter failure.

#### 2.4.4 Drug Self-administration – Maintenance

After the 1^st^ saline-substitution session (see below), rats continued under the same T_A_ conditions (and priming criterion) as in acquisition phase for an additional 11 sessions. A 2^nd^ saline substitution immediately followed this Maintenance phase (see below).

#### 2.4.5 Drug Self-administration – Temperature-Condition Swap

After the 2^nd^ saline substitution (see below), rats continued under the same T_A_ conditions and priming criterion as in maintenance for one additional IVSA session (the 30th following group assignment; 33rd overall) after which the groups’ T_A_ conditions were swapped. Switched T_A_ conditions continued for seventeen sessions during which a 3^rd^ saline substitution was intercalated just prior to the final four sessions. Five rats from the group that begin self-administration under the 30 °C T_A_ condition were unable to complete this phase (4 due to loss of patency and 1 due to poor health requiring euthanasia per protocol), inclusive of the two that did not complete the second saline substitution, see below. All of the rats that begin self-administration under the 20 °C T_A_ condition were able to complete this phase. Thus, in this phase of the experiment, 6 Hot-trained rats changed from a 30 °C to 20 °C T_A_ and 12 Cold-trained rats changed from a 20 °C to 30 °C T_A_.

#### 2.4.6 Saline Substitutions

Three sessions were given wherein MDMA was replaced with saline to probe behavior and body temperature responses at various points in the series of IVSA sessions. The first of these was given within a single session at the end of the Acquisition phase (16th from start of altered temperature conditions, 19th overall). In this case the number of infusions during this saline day were compared with infusions on the prior and succeeding days on which MDMA was available. Unfortunately, a spike in room temperature occurred on this saline day in the “Hot” room (∼2 °C above the prior and subsequent MDMA sessions). Thus, the latter two saline-substitution sessions were each conducted over two sequential sessions wherein MDMA or saline were available on one of those two sessions in counterbalanced order to account for any variance between days. These were sessions 28-29 for the 2^nd^ substitution (the end of the Maintenance phase) and sessions 44-45 for the 3^rd^ substitution (near the end of the Temperature Swap phase). Two rats from the “Hot-trained” group were excluded from the 2^nd^ substitution (one due to loss of patency; the other due to exceeding a body temperature of 40 °C – it self-administered 15 mg/kg of MDMA at a rate of 22 mg/kg/hr on that particular session). Exclusions for the 3^rd^ saline substitution were the same as those for the Temperature Swap phase (detailed above). Saline substitutions were done under whichever T_A_ condition the group was training under at the time.

#### 2.4.7 Non-contingent Drug Administration

After the final temperature condition swap session, rats were given 10 sessions wherein MDMA was delivered i.v. non-contingently under a consensus timing pattern derived from the average intervals derived from the self-administration sessions. The T_A_ was either 20 °C or 30 °C (order counterbalanced) and the per-session dose ranged from 1 mg/kg to 5 mg/kg (in an escalating dose order). Infusions began after a 15-min baseline period and were spaced by the following formula: Inter-infusion Interval (sec) = -17*X^2^ + 287*X - 338; where X is the infusion number. This curve approximates the pattern of inter-infusion intervals observed in those rats from this experiment that averaged eight or more infusions per session. One of the rats from the group that begin self-administration under the 30 °C T_A_ condition and one of the rats from the group that begin self-administration under the 20 °C T_A_ condition were unable to complete this phase (both due to loss of catheter patency). Thus, this phase of the experiment included 5 Hot-trained and 11 Cold-trained rats. By this time, the Hot-trained rats had a total of 28 MDMA IVSA sessions at a T_A_ of 30 °C and 18 sessions at a T_A_ of 20 °C (not including saline sessions); *vice versa* for the Cold-trained rats.

### 2.5 Experiment 2: Self-stimulation reward

#### ICSS Procedure

Rats were tested in sound attenuating operant chambers, which contain a wheel manipulandum within one wall. Trials begin with a noncontingent stimulation (sinusoidal electrical stimuli of 250ms duration and 60Hz), followed by a variable post-stimulation interval (7.5 s) during which delivery of a second stimulus was contingent upon responding with a 1/4 turn of a wheel manipulandum. Each electrical stimulation (reinforcer) had a train duration of 500 ms during which 0.1 ms cathodal pulses were delivered at 50-100 Hz, with current-intensity thresholds within 50-200μA. Current was varied in a series of steps (+5μA per step, 3 trials per step). In each testing session, four alternating descending-ascending series were presented. The threshold for each series was defined as the midpoint between two consecutive current intensities that yielded ‘positive scores’ (animals responded for at least two of the three trials) and two consecutive current intensities that yielded ‘negative scores’ (animals did not respond for two or more of the three trials). The overall threshold of the session was defined as the mean of the thresholds for the four individual series. Each testing session was ∼30 min in duration. Rats were trained once daily until stable reward threshold were established (≤10% variation in thresholds for three consecutive days) between 7 and 10 days.

To determine the effect of MDMA and ambient temperature, animals were tested twice per experimental day with the first determination at 24°C, which served as the baseline. Two hours later thresholds were re-determined in 18°C or 30°C ambient temperature conditions and expressed as a percent of the individual’s baseline threshold for that day. Animals were introduced to the testing temperature 30 min prior to the second session and injected (s.c.) 10 min prior to the start of the second session with either 2.5 mg/kg MDMA or saline vehicle. The order of the four conditions was randomized across individuals.

### 2.6 Designs and Data Analyses

IVSA measures of drug intake (total infusions per 2-hour session), lever discrimination (% correct), body weight, body temperature (°C), and activity rate (counts/min) were analyzed as functions of the between-subjects factor of T_A_ group (Hot-trained or Cold-trained) and within-subjects factors of session. Body temperature and activity rates during IVSA were analyzed as function of T_A_ group and the within-subjects factors of block of sessions (3 session/block for baseline, 5-6 sessions/block for all subsequent phases) and time from lever extension (one pre-session reading followed by binned 30-min time intervals). The non-contingent drug administration measures of body temperature and activity were analyzed as a function of the between subjects factor of MDMA-self-administration grouping (based on mean intake of the last 4 self-admin sessions; 0-2 mg/kg vs. 4-10 mg/kg) and the within-subjects factors of ambient temperature (20 °C vs. 30 °C), MDMA dose (1-5 mg/kg/session; delivered in a pattern based on self-administration) and time from session start (one 15-min pre-session binned time interval followed by three 60-min binned time intervals). Intracranial self-stimulation reward thresholds (% of baseline) were analyzed as a function of the within-subjects factors of drug treatment (MDMA vs Vehicle) and T_A_. All analyses were performed using repeated-measures analysis of variance (rmANOVA). Effects confirmed by rmANOVA were delineated with post hoc comparisons using Tukey’s HSD (IVSA) or Neuman-Keuls (ICSS).

StatView (SAS Institute, Inc., Cary, NC) was used for IVSA analyses and GB-STATv7.0 (Dynamic Microsystems, Silver Spring MD) was used for ICSS analyses. Graphs were generated with Microsoft Excel (Microsoft, Redmond WA) and StatView and figures created with Canvas (ACD Systems, Seattle WA).

## 3. Results

### 3.1 Experiment 1: Drug Self-Administration under Varied Ambient Temperature (T_A_)

#### 3.1.1 Drug Self-administration – Baseline

##### 3.1.1.1 Infusions

There was a progressive reduction in MDMA intake (**Figure 1**) across the three baseline sessions (T_A_ = 25° C for all) as confirmed by a main effect of session (F_2,42_= 15.01; p<0.001). The post-hoc analysis likewise confirmed that significantly fewer infusions were obtained in the second and third sessions relative to the first session. Groups were successfully balanced on intake (i.e., there was no confirmed effect of group). Similarly, lever discrimination increased from the first session (Mean = 83%; SEM=±4.5%) to the second session (94%, ±1.7%), but neither first nor second sessions were different from the 3^rd^ session (91%, ±2.2%) as confirmed by a main effect of session on lever discrimination (F_(2,42)_ = 3.52; p < 0.05), but no main effect of (future) T_A-_training group or interaction between factors.

**Figure 1:**
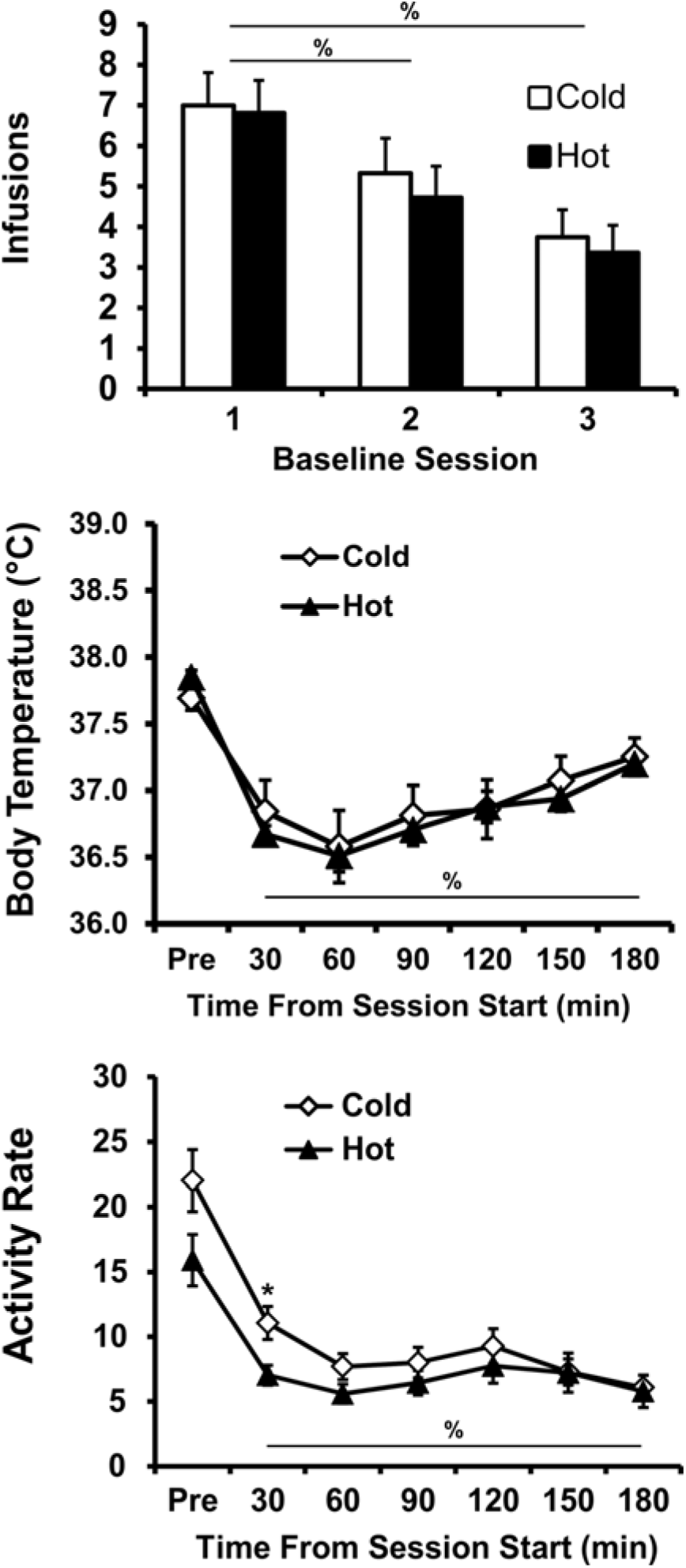
Mean (±SEM) MDMA infusions, body temperature (°Celsius) and activity rate (counts/min) during the three baselining sessions. Data are grouped by the subsequent intake-balanced assignments to either the Cold (20° C; N= 12) or Hot (30° C; N= 11) ambient temperature training conditions. Means significantly different from the first session (infusions) or pre-session baseline (body temp, activity) are indicated by % and differences across groups by *.

##### 3.1.1.2 Temperature

The body temperature of the rats (Figure 1), averaged across the three baseline days, declined significantly across the IVSA session (F_6,126_= 59.86; p<0.0001) but there was no difference between the groups. Post hoc comparisons confirmed that body temperature was lower than the baseline value at all subsequent timepoints (30-180 min after the start of the session).

##### 3.1.1.3 Locomotor Activity

Activity (**Figure 1**), averaged across the three baseline days, also declined significantly within session (F_(6,126)_ = 50.74; p < 0.0001) and the future T_A_-training groups did differ in baseline activity as confirmed by an interaction between group and time bin (F_(6,126)_ = 2.91; p = 0.01), although there was no main effect of future T_A_-training group. Post hoc comparisons confirmed that pre-session activity was higher than that of any other time for each group, and activity was greater in the first 30 min than in the third hour for the future train-in-cold group only. Activity differed between groups in the first 30 min only. To rule out the possibility that these activity differences could predict subsequent MDMA self-administration intake, correlational analyses (Fisher’s z-tests) were conducted between end-of-acquisition infusions and activity in the first 30 min of baseline (the only time bin with a reliable between-group difference). Activity and number of infusions did not correlate significantly either as a combined group (r^2^ = +0.001, Z_(23)_ = 0.15, p = 0.88) or as separate groups (start-in-hot, r^2^ = +0.092, Z_(11)_ = 0.89; p = 0.37; start-in-cold, r^2^ = +0.261, Z_(12)_ = 1.69; p = 0.09).

Thus, this difference in activity between the future T_A_-training groups did not predict end-of-acquisition intake values.

#### 3.1.2 Drug Self-administration – Acquisition

##### 3.1.2.1 Infusions

In the acquisition phase, animals continued IVSA, but under either a Hot (30°C; N=11) or a Cold (20°C; N=12) ambient temperature (T_A_). Intake increased for both group, but this increase was greater for the Hot group (**Figure 2**). Analysis of drug intake confirmed a main effect of session (F_14,294_=6.09; p<0.001) and the interaction of session with T_A_ group (F_14,294_= 2.29; p<0.01) but no main effect of group.Post-hoc comparisons confirmed that intake was greater than that of the first acquisition session in sessions 13-15 for the Hot group and 6 and 13 for the Cold group. Furthermore,intake was greater in the Hot group than the Cold group on the final (15^th^) session. Lever discrimination remained high and consistent for both the Hot (83%, ± 3.2%) and Cold (84%, ± 4.6%) groups. Analysis of lever discrimination confirmed that there was no main effect of T_A_ group or session, nor was there an interaction between these factors.

**Figure 2:**
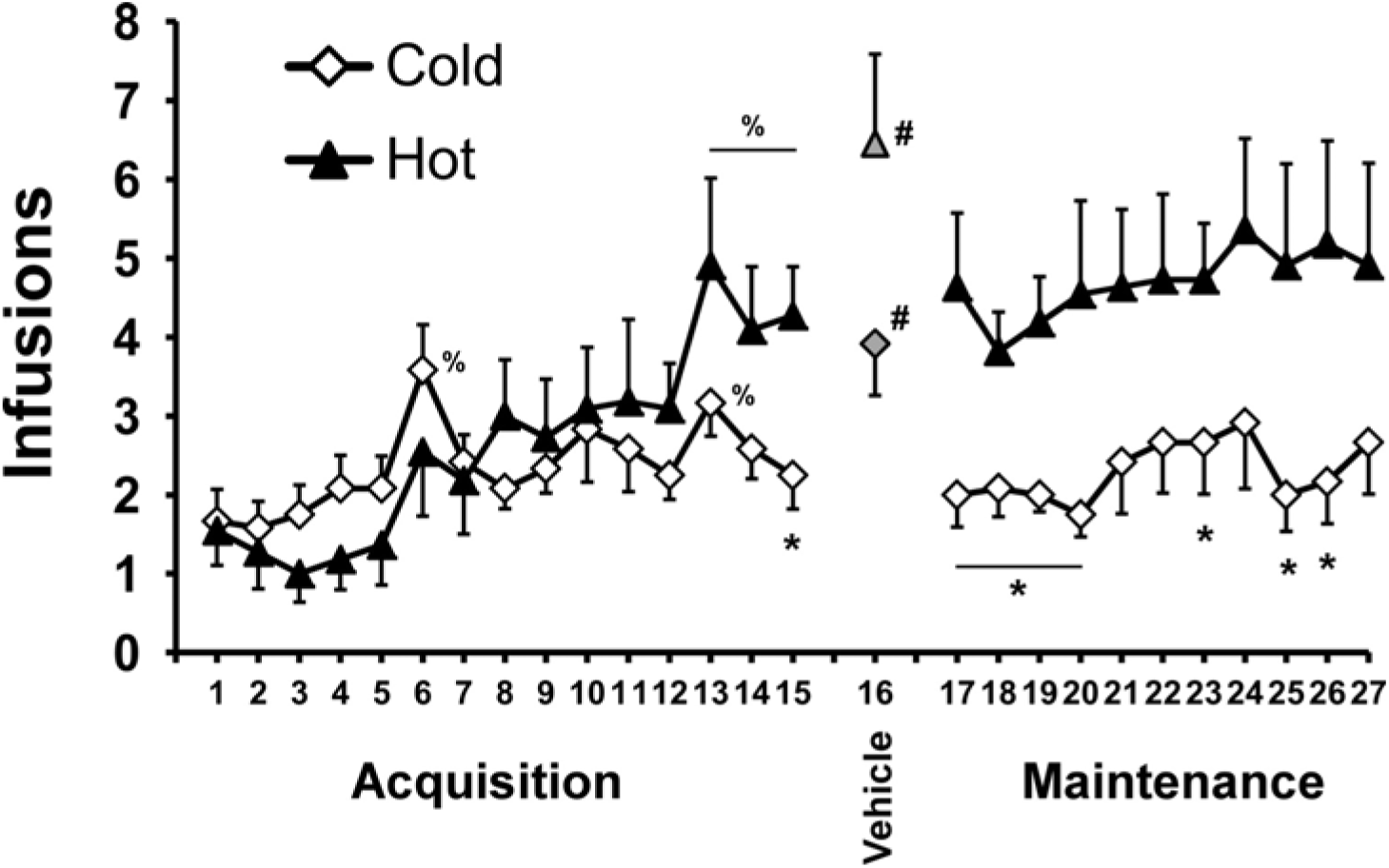
Mean (±SEM) infusions of MDMA as a function of session of Acquisition (sessions 1-15), Maintenance (sessions 17-27) and the 1^st^ saline substitution (session 16) grouped by the ambient temperature (Hot = 30° C, Cold = 20° C) under which self-administration occurred. Significant differences between group is indicated by *, differences from first session (except saline) by % and between the saline substitution and the average of the prior and subsequent sessions by #.

##### 3.1.2.2 Temperature

The body temperature of rats continued to exhibit a characteristic drug-associated decline throughout the Acquisition and Maintenance intervals (**Figure 3**). The data for the Acquisition was grouped in 5-session blocks thus the analysis included a between subjects factor of Group and within-subjects’ factors of block and time within the session. The ANOVA confirmed a main effect of block (F_2,42_= 37.24; p<0.0001), time within the session (F_6,126_= 86.46; p<0.0001) and the interaction of these two factors (F_12,252_= 2.60; p<0.005). The analysis also confirmed a significant interaction of group with block (F_2,42_=3.76; p<0.05) and the interaction of block with time (F_12,252_= 3.30; p<0.0005).

**Figure 3:**
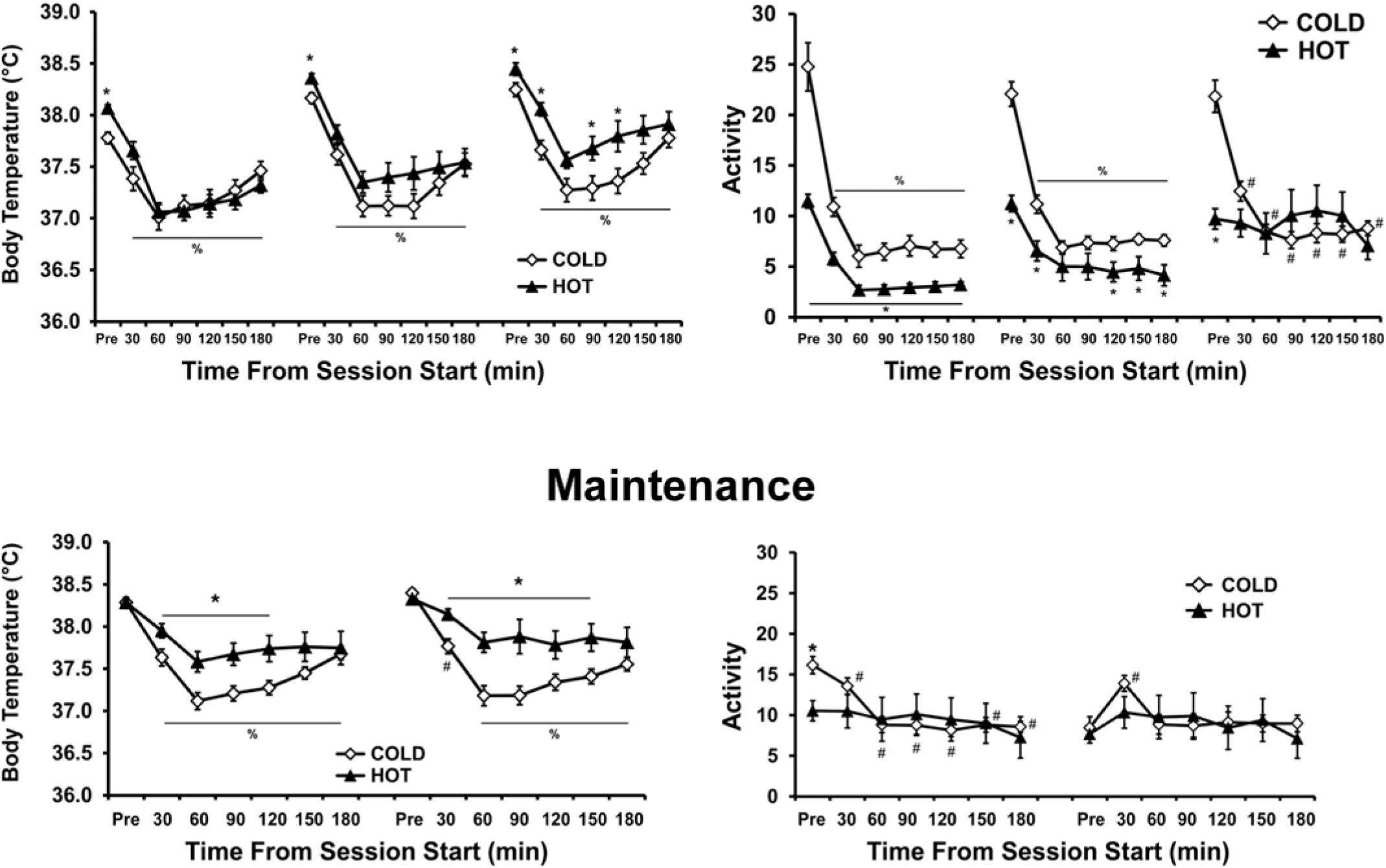
Mean (±SEM) body temperature and activity rate of the groups trained under either a Hot (30° C) or Cold (20° C) ambient temperature, as a function of time from the start of sessions (one pre-session time point followed by binned 30-min time intervals; drug available only for the 1^st^ 2 hours). Data from individual self-administration sessions are collapsed across blocks of 5-6 sessions during Acquisition (top) and Maintenance (bottom) intervals. A significant difference between groups is indicated with *, differences from the respective pre-session temperature (Pre) for both groups by %, and differences from the Pre value for the Cold group only by #.

The post-hoc analysis confirmed first that body temperature was lower for all time points (30-180 min after session start) compared with the pre-session baseline for both groups in each of the three blocks. The post-hoc analysis also confirmed that baseline temperatures for the Hot group were higher than that of the Cold group for all three blocks. There were no between-group differences confirmed for any time point after the session start for the first two blocks. However, body temperature was significantly lower in the Cold group for the 30, 90 and 120 min time points in the final block. Also, for the Hot group, body temp increased for each bin in each subsequent block with exception of the pre-session bins of the middle and final blocks being equivalent. Lastly, for the Cold group, although body temperatures also increased with each subsequent block, the pattern was less consistent (initial < middle for Pre & 30 min, initial < final for Pre to 60 min and 150-180 min, middle < final for 120 and 180 min). Thus, body temperature increased across acquisition for both groups, but this increase was larger and more consistent across time bins in the Hot group.

##### 3.1.2.3 Locomotor Activity

Analysis of activity (**Figure 3**) confirmed main effects of group (F_(1,21)_ = 9.84; p < 0.01), block (F_(2,42)_ = 9.86; p < 0.001) and time within the session (F_(6,126)_ = 109.31; p < 0.0001), as well as interactions between group and block (F_(2,42)_ = 4.08; p < 0.05), Group and time (F_(6,126)_ = 27.29; p < 0.0001), block and time (F_(12,252)_ = 9.09; p < 0.0001), and all three factors (F_(12,252)_ = 2.81; p < 0.01). Post hoc comparisons confirm that activity was lower in the Hot group than the Cold group for all time points in the initial block of sessions, for the all but the 30-min and 60-min time bins in the middle block and only in the pre-session sample in the final block. Within the Hot group, activity was higher pre-session than in all subsequent time bins for the initial and middle blocks – but not for the final block – and activity was higher in the first 30-min time bin than all subsequent time bins in the initial block. For the Cold group activity was higher pre-session than all other time bins and higher in the first 30-min bin than all subsequent time bins in all 3 blocks. Finally, activity was higher for the Hot group in the final block compared with the first two blocks for all time bins of the last two hours of the session, as well as higher than the initial block in the first hour of the session. In contrast, activity for the Cold group was only higher in the final block compared with the initial block for the 30-60 min and 150-180 min time bins. Thus, a higher ambient temperature was associated with lower initial activity levels during MDMA self-administration, but that T_A_ effect diminished as the activity of the hot-T_A_ group increased in each subsequent block of sessions.

#### 3.1.3 Drug Self-Administration – Maintenance

##### 3.1.3.1 Infusions

The difference in MDMA self-administration between Hot (N=11) and Cold (N=12) groups which emerged at the end of the acquisition interval was sustained during the 11 session maintenance interval, although intake was relatively stable across sessions within each group (**Figure 2**). The analysis of the maintenance phase confirmed main effects of group (F_1,21_= 6.45; p<0.05), but not of session, nor was there a significant interaction between the factors. Post-hoc comparisons confirmed significant group differences for sessions 17-20, 23 and 25-26. As in Acquisition, lever discrimination in the Maintenance phase remained stable and similar for both the Hot group (84%, ± 4.1%) and the Cold group (84%, ± 4.8%). Analysis of lever discrimination confirmed that there was no main effect of T_A_ group or session, nor was there an interaction between these factors.

##### 3.1.3.2 Temperature

The body temperature data was binned and pooled into two 5-to-6-session blocks. Analysis of body temperature during Maintenance (**Figure 3**) confirmed main effects of group (F_1,21_=7.80; p<0.05) and time within the session (F_6,126_= 43.41; p<0.0001), as well as the interaction between group and time (F_6,126_=5.43; p<0.0001). Post hoc comparisons confirmed that body temperature was lower after the session start in the Cold group compared with the Hot group 30-120 min the first half of Maintenance and 30-150 min in the second half of the Maintenance phase. Similarly, all body temperatures were lower than the pre-session values except for that of the 30 min time bin for the Hot group in the second half of Maintenance.

##### 3.1.3.3 Locomotor Activity

Analysis of activity confirmed main effects of time within-session (F_(6,126)_ = 6.07; p < 0.0001) – but, not group or session block – as well as significant interactions between group and time (F_(6,126)_ = 42.14; p < 0.01), block and time (F_(6,126)_ = 47.30; p < 0.0001) and all three factors (F_(6,126)_ = 13.15; p < 0.01). Post hoc comparisons confirmed that the Hot group had higher activity than the Cold group only pre-session in the first block. There were no significant activity differences between any time-bin in the Hot group for either block. Activity for the Cold group was *higher* pre-session than in subsequent time bins in the first block and *lower* pre-session compared with the first 30-min bin in the second block. Additionally, Cold-group pre-session activity was higher than in subsequent bins for both blocks. Lastly, regardless of T_A_ group, activity was higher in the first than in the second block for pre-session activity only.

#### 3.1.4 Drug Self-administration – Temperature-Condition Swap

##### 3.1.4.1 Infusions

Switching animals between ambient temperature conditions did not change the group difference in drug intake; rather, the Hot-trained group (N=6), now in a cold T_A_, continued to increase intake while the Cold-trained group (N=12), now in a hot T_A_, largely maintained intake levels but did not drop below initial intake levels on 6 of the 17 sessions (**Figure 4**). The ANOVA confirmed main effects of group (F_1,16_= 6.47; p<0.05) and session (F_17,272_ = 2.67; p<0.0005), as well as the interaction of group with session (F_17,272_ = 3.17; p<0.0001). Post-hoc comparisons confirmed that more infusions were obtained by the Hot-trained group in sessions 41-43 and 46-49. Additionally, relative to the session prior to the temperature swap, the Cold-trained group obtained *fewer* infusions in sessions 36, 39-42 and 46 while the Hot-trained group self-administered *more* infusions in sessions 47 and 49. As in the prior two phases,overall lever discrimination in the Temperature-Swap phase remained stable and similar for both the Hot-trained group (89%, ± =4.7%) and the Cold-trained (87%, ± 2.8%) groups. The analysis of lever discrimination confirmed that there were no significant effects of T_A_ group, of session, or of the interaction of factors.

**Figure 4:**
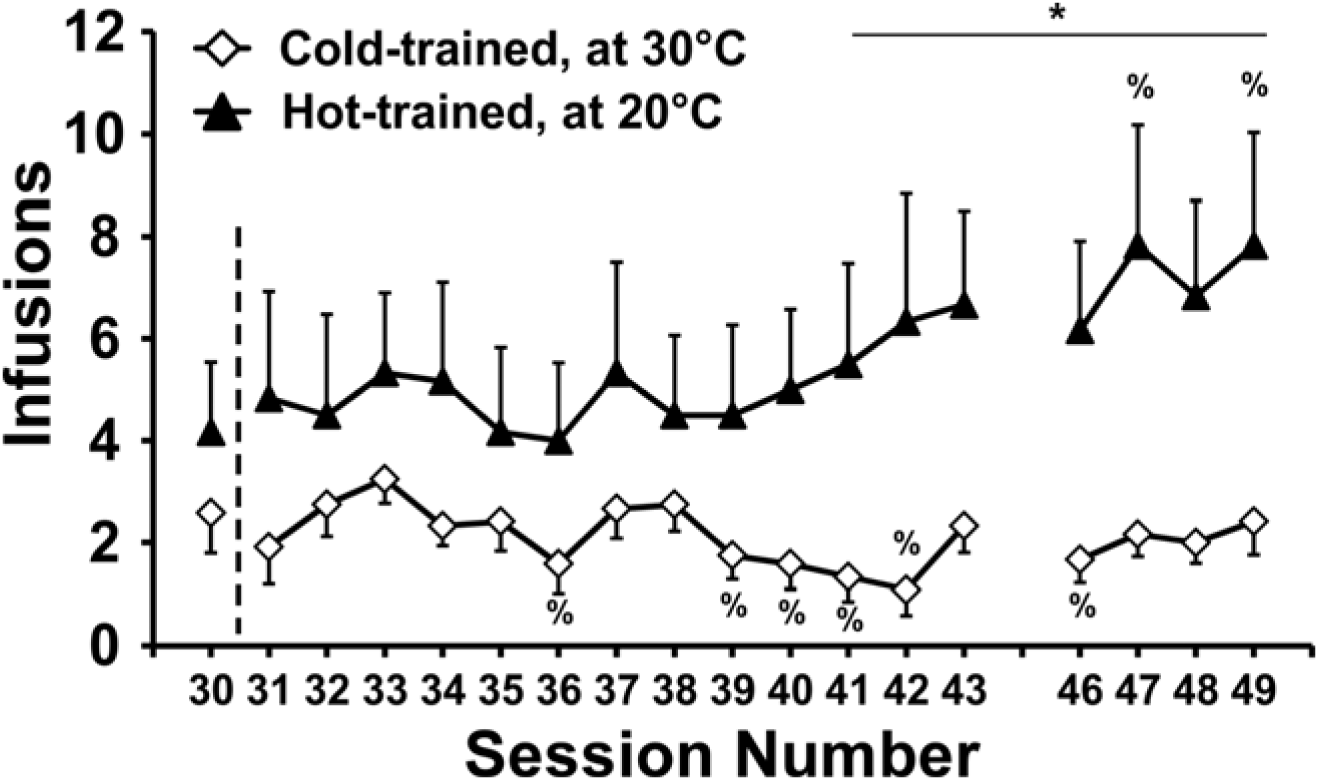
Mean (±SEM) infusions of MDMA as a function of sessions either before (session 30) or after swapping the ambient temperature (T_A_) conditions for the groups. Thus, for sessions 31-49, *Hot-trained* animals were tested under *20° C* and *Cold-trained* animals were tested under *30° C*. Missing sessions (44-45) were those of the 3^rd^ saline substitution, reported in the text. Significant differences between groups are indicated by * and differences from the pre-swap session (30) within-group are indicated by %.

##### 3.1.4.2 Temperature

The body temperature data were again binned and blocked similar to that for preceding phases (5-6 sessions per block). Body temperature decreased over time within sessions (**Figure 5**) and was overall lower in the Hot-trained group than the Cold-trained group for each block (as expected from the T_A_-swap). The ANOVA confirmed main effects of Group (F_1,15_= 14.28; p<0.005), session block (F_2,30_= 11.76; p<0.0005) and time within-session (F_6,90_= 134.38; p<0.0001) as well as the interactions of Group with time (F_6,90_= 4.40; p<0.005) and of block with time (F_12,180_= 2.09; p<0.05). Post hoc comparisons confirmed significant differences between the groups for time bins in the initial (Pre-90 min), middle (30-150 min) and final (Pre-180) blocks of sessions. Body temperature in both groups declined significantly within sessions as compared with pre-session values for all time bins and session blocks. Body temperature for the Hot-trained group differed between initial and final session blocks only in the first 30 min. In contrast, body temperature for the Cold-trained group was higher in the final block compared with the initial block for all but the pre-session bin, higher in the final block than middle block for the first 2 hours, and higher in the middle block than first block in the last hour of the session.

**Figure 5:**
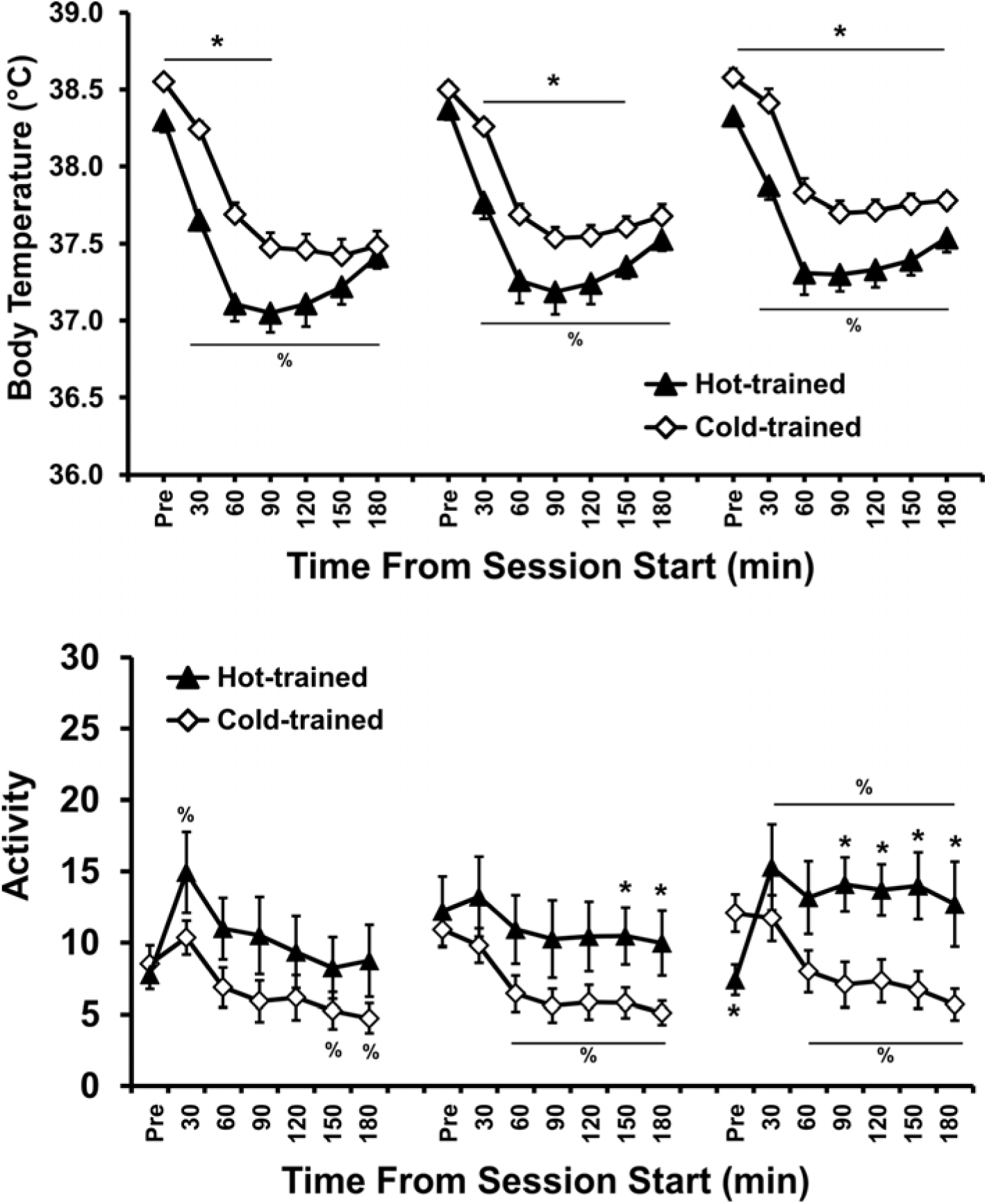
Mean (±SEM) body temperature and activity rate for the Hot-trained or Cold-trained groups during the Temperature-Swap phase (i.e., *Hot-trained* animals were tested under *20° C* and *Cold-trained* animals were tested under *30° C*) as a function of time from the start of sessions and block of sessions (5-6 sessions per block). Significant differences between treatment groups is indicated by * and within-group differences from the pre-session value are indicated by %.

##### 3.1.4.3 Locomotor Activity

The analysis of activity (**Figure 5**) confirmed main effects of session block (F_(2,34)_ = 14.66; p < 0.0001) and time within-session (F_(6,102)_ = 12.00; p < 0.0001) – but, not group – as well as significant interactions between group and time (F_(6,102)_ = 7.30; p < 0.0001), block and time (F_(12,204)_ = 5.54; p < 0.0001) and all three factors (F_(12,204)_ = 4.79; p < 0.0001).

Post hoc comparisons confirmed that activity was higher in the Hot-trained group than the Cold-trained group in the last hour of the middle block and the last 2 hours of the final block, and pre-session activity was lower in the Hot-trained group than the Cold-trained group in the final block. For the Hot-trained group, activity *increased* above pre-session levels for only the 1^st^ 30 min of the session in the initial block, for none of the bins in the middle block, but for the entire session in the final block. However, for the Cold-trained group, activity *decreased* below pre-session levels in the last hour of the initial block, the last 2.5 hrs of the middle and final blocks.

Finally, activity was lower for the Hot-trained group in the initial block compared with the final block for the last 1.5 h of the session and higher pre-session in the middle block compared with both the first and final blocks. For the Cold-trained group, activity was lower pre-session in the initial block compared with the last two blocks, higher in the last block than the middle block in the first hour of the session, and higher in the final block than the initial block in the 120-150 min bins. Thus, as in the acquisition phase, activity during the temp-swap phase was initially overall comparatively lower under the high T_A_ condition. However, in this temp-swap phase, the activity differences between the T_A_ conditions overall widened across blocks while activity differences all but disappeared by the end of the maintenance phase (i.e., the block of sessions just prior to reversing the T_A_ conditions).

#### 3.1.5 Saline Substitutions

##### 3.1.5.1 Infusions

The first saline substitution session was run immediately after the Acquisition phase (i.e., the 16^th^ session in **Figure 2**) in the Hot (N=11) and Cold (N=12) groups. The room temperature was inadvertently higher than the target 30 (±1)°C for the Hot (N=11) group, thus body temperature was not analyzed for this first substitution and conclusions drawn from the analysis of infusions must be tempered. This analysis compared infusions on the saline day with the prior session (final day of the acquisition phase) and the subsequent session (first day of the maintenance phase). The rmANOVA confirmed main effects of both T_A_ Group (F_1,21_= 8.62; p<0.01) and session (F_2,42_= 8.19; p<0.001), but no interaction. Post-hoc comparisons further confirmed that infusions were significantly higher during the saline day than during either MDMA session (which did not differ from each other). In addition, the number of infusions was higher in the Hot group than in the Cold group on both MDMA sessions but not on the saline session.

Similar results for the analyses of infusions were obtained for the 2^nd^ and 3^rd^ saline substitutions. In these analyses, saline or MDMA were available, in counterbalanced order, on two consecutive sessions. In the second saline substitution conducted at the end of the maintenance phase, the Cold-trained group (N=12) obtained 2.5 (SEM: 0.3) MDMA infusions and 5.9 (1.2) saline infusions. The Hot-trained group (N=9) obtained 5.8 (1.3) MDMA infusions and 7.6 (1.3) saline infusions. The ANOVA confirmed main effects of T_A_ group (F_1,20_ = 4.76; p < 0.05) and Substitution (F_1,20_ = 6.36; p < 0.05), but no interaction (F_1,20_ = 0.61; p = 0.44). Similarly, for the third substitution, the Cold-trained group (N=12) obtained 2.0 (SEM: 0.6) MDMA infusions and 3.7 (0.8) saline infusions and the Hot-trained group (N=6) obtained 4.8 (1.7) MDMA infusions and 8.2 (2.2) saline infusions. The rmANOVA confirmed main effects of T_A_ group (F_1,16_ = 6.07; p < 0.05) and Substitution (F_1,16_ = 10.96; p < 0.01), but not of the interaction. Post hoc comparisons confirmed that in both substitutions, more infusions were earned when saline was available than when MDMA was available and more infusions were earned by the Hot-trained group than the Cold-trained group. Lever Discrimination remained stable across conditions and comparable between groups for all three saline substitutions (analyses confirmed no significant main effects or interactions; all p > 0.4).

##### 3.1.5.2 Temperature

Body temperature was differentially affected by MDMA and saline IVSA for both the second and third saline substitutions (**Figure 6**). Overall, body temperature dropped below that of saline under a T_A_ 20° C, but not 30° C, regardless of prior T_A_-training. Analysis of body temperature for the second substitution confirmed a significant main effect of time within-session (F_(6,114)_ = 19.731; p < 0.0001) as well as interactions between T_A_ Group and time (F_(6,114)_ = 2.905; p < 0.05), drug condition and time (F_(6,114)_ = 2.959; p < 0.05), and all three factors (F_(6,114)_ = 3.493; p < 0.01). There were no main effects of T_A_ Group or drug condition confirmed. Post hoc comparisons for the Cold-trained group confirmed that body temperature declined significantly from pre-session baseline values for all time bins for both MDMA and saline sessions. For the Hot-trained group, body temperature decreased from baseline from 60-180 min for the saline session, but only in the 60-min time bin for the MDMA session. Additionally, for the Cold-trained group, body temperature was lower for the MDMA session than the saline session for the 30and 90-180-min time bins. However, for the Hot-trained group, significant differences between saline and MDMA sessions were not confirmed.

**Figure 6:**
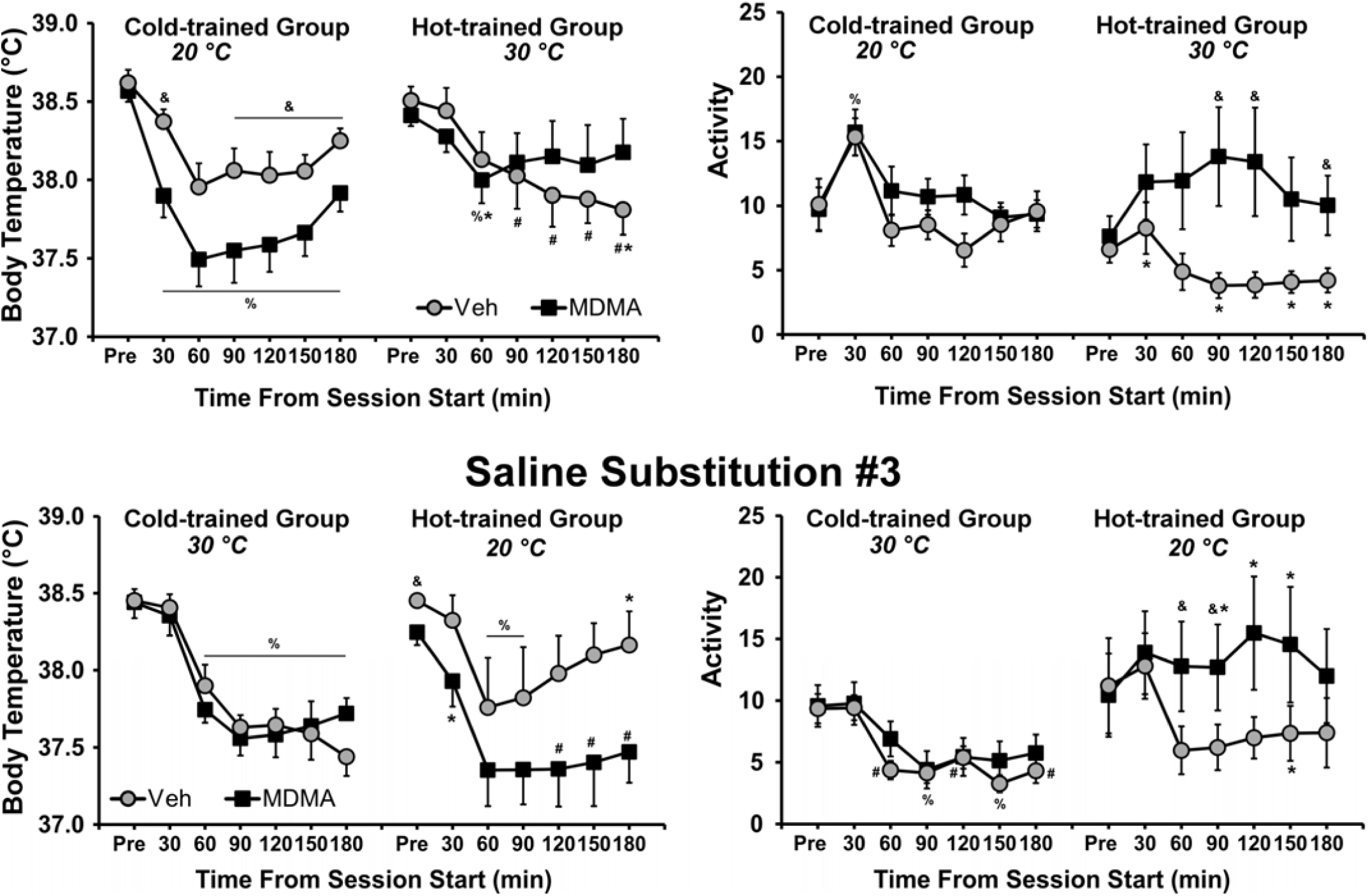
Mean (±SEM) body temperature and activity rate during the second and third saline substitution experiments are presented for Hot-trained or Cold-trained groups as a function of time from the start of sessions and saline/MDMA drug availability. A significant difference between drug treatment conditions, within group, is indicated with & and differences between groups, within drug treatment by *. Significant differences from the respective pre-session body temperature (Pre) for both treatment conditions are indicated by %, and differences from the Pre value for one treatment condition by #.

Analysis of body temperature for the third saline substitution (under the T_A_-condition swap)confirmed significant main effects of drug condition (F_(1,12)_ = 5.44; p < 0.05) and time (F_(6,72)_ = 28.51; p <0.0001) and as well as interactions between T_A_-trained group and time (F_(6,72)_ = 2.38; p < 0.05) and T_A_-trained group and drug condition (F_(1,12)_ = 5.3; p < 0.05). There was no significant effect of group. Post hoc comparisons for the Cold-trained group confirmed that body temperature declined significantly from pre-session baseline values for all post-injection time points for both MDMA and saline sessions. For the Hot-trained group, body temperature decreased from baseline from 60-180 min for the MDMA session, but only from 60-90 min for the saline session. Additionally, for the Hot-trained group, body temperature was lower for the MDMA session than the saline session only in the pre-session baseline. However, for the Cold-trained group, there were no significant differences between saline and MDMA sessions.

##### 3.1.5.3 Locomotor Activity

Activity was also affected by treatment and training group. Analysis of activity for the second substitution (**Figure 6**) confirmed significant main effects of drug condition (F_(1,19)_ = 10.59;p < 0.01) and time (F_(6,114)_ = 6.08; p < 0.0001) as well as a significant interaction between drug condition and Time (F_(6,114)_ = 4.44; p < 0.001) and a trend toward an interaction between T_A_ and drug condition (F_(1,19)_ = 4.22; p = 0.05). There was no main effect of T_A_ nor of the three-way interaction confirmed. Post hoc comparisons confirmed that compared to saline, activity was higher in the Hot-trained group when MDMA was available (under the 30° C-T_A_ condition) 90, 120 and 180 min after the session start while there were no drug-related differences in Cold-trained group activity. Also, activity was lower in the Hot-trained group compared with the Cold-trained group when saline was available (30, 90, 150, 180 min). Within-session activity only changed for the Hot-trained group when saline was available, and activity was higher in the first 30-min time bin compared with the last 2 hours. Activity for the Cold-trained group was higher in the first 30-min time bin than at all other times, when either saline or MDMA was available.

Analysis of activity for the third substitution (*under the T*_*A*_*-condition swap*) confirmed significant main effects of T_A_-condition (F_(1,16)_ = 6.21; p < 0.05), drug condition (F_(1,16)_ = 5.26; p < 0.05) and time within-session (F_(6,96)_ = 3.98; p < 0.01) as well as a significant interaction between drug condition and time (F_(6,96)_ = 2.76; p < 0.05); the three-way interaction was not significant (F_(6,96)_ = 1.90; p = 0.09). Post hoc comparisons confirmed that activity was higher when MDMA was available, compared to saline, in the original Hot-trained group (now run in the 20° C-T_A_) 60-90 min after session start, but there were no activity differences between saline and MDMA for the original Cold-trained group (now run in the 30° C-T_A_). Activity was higher in the Hot-trained group compared with the Cold-trained group when MDMA (90-150 min after session start) or saline (150 min) was available. Activity did not vary across time for the Hot-trained group, but did decline relative to the pre-treatment baseline in the Cold-trained group when saline (60-180 min after session start) or MDMA (90, 150 min) was available.

#### 3.1.6 Non-contingent Drug Administration

##### 3.1.6.1 Temperature

The animals were divided into high (N=5) and low (N=10) MDMA exposure groups based on the average amount of MDMA self-administered (0-2.0 mg/kg vs 3.5-12 mg/kg; High-SA vs. Low-SA) in the four days preceding the non-contingent sessions, rather than by training group. MDMA differentially reduced body temperature in a manner dependent upon the non-contingent dose given, and the self-determined prior dose-range grouping, as is shown in **Figure 7**. The main analysis of body temperature included a between-subjects factor of self-administration Group and the within-subjects’ factors of time (Baseline, 1h, 2h, 3h), ambient temperature (T_A_; 20°C vs 30°C) and the non-contingent MDMA dose (1-5 mg/kg). The ANOVA confirmed that body temperature was significantly affected by ambient temperature (T_A_) (F_1,13_=31.047; p<0.0001), MDMA Dose (F_4,52_=13.376; p<0.0001) and time (F_3,39_=104.174; p<0.0001). Furthermore, the analysis confirmed significant interactions between T_A_ and dose (F_4,52_=15.56; p<0.0001), T_A_ and time (F_3,39_=30.91; p<0.0001), dose and time (F_12,156_=9.269; p<0.0001) and of all three of these factors (F_12,156_=10.608; p<0.0001). Although the main effect of group did not reach statistical significance, there was a significant group X time interaction (F_3,39_=6.961; p<0.001) and group X dose X time interaction (F_12,156_=2.112; p<0.05).

**Figure 7:**
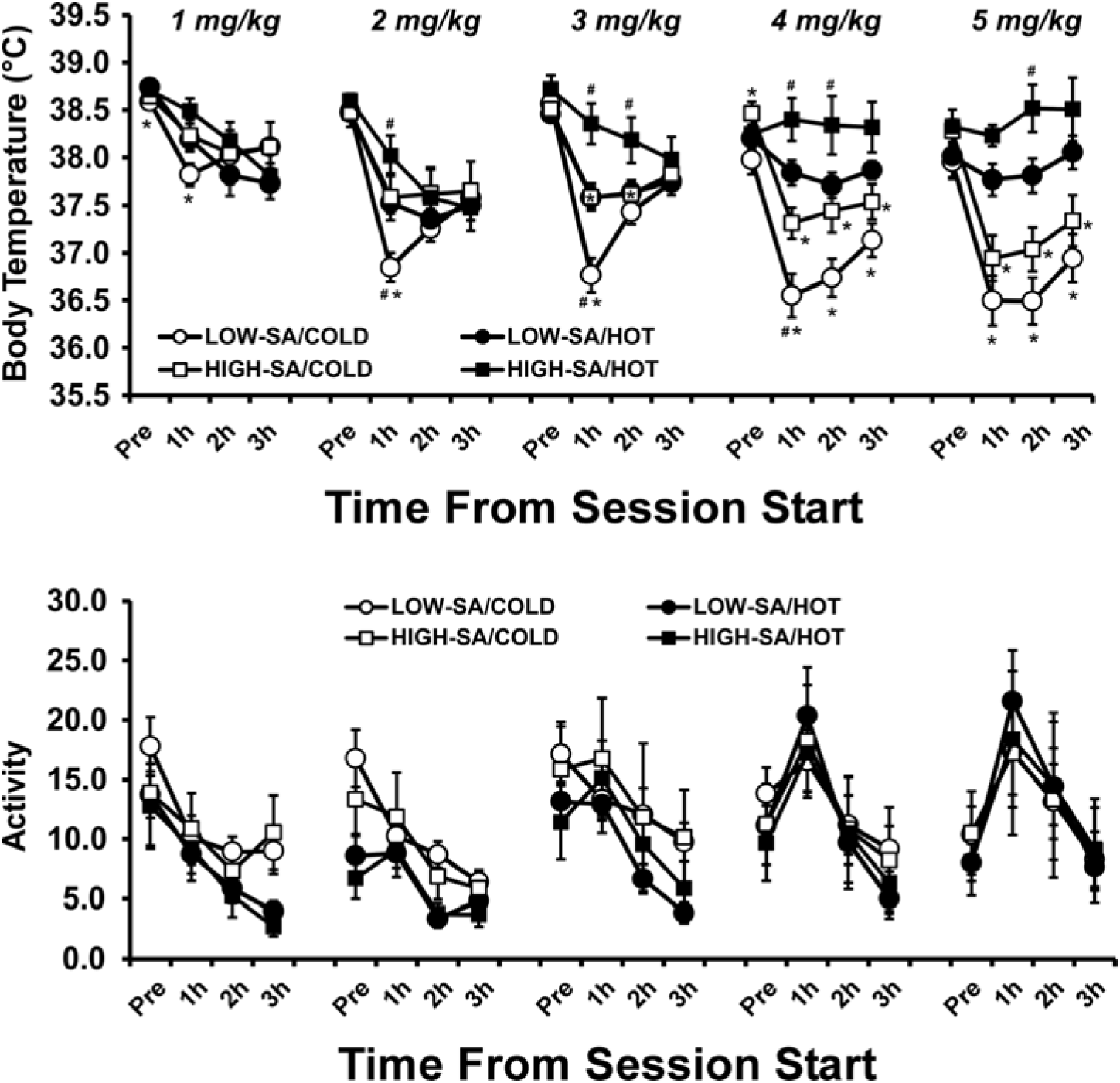
Mean (±SEM) body temperature and activity rates following the non-contingent i.v. administration of MDMA is presented for two groups, determined by the average number of infusions obtained over the final four self-administration sessions (LOW-SA = 0-2.0 mg/kg; HIGH-SA = 3.5-12 mg/kg). Each group was challenged under two ambient temperatures (HOT = 30° C, COLD = 20° C; order balanced across dose) and five total MDMA doses per session (1, 2, 3, 4 or 5 mg/kg; given in ascending order). Significant differences from the pre-session baseline (Pre) are indicated by %, differences between groups are indicated by * and differences between ambient temperature conditions, within group, are indicated by #.

The post hoc comparisons confirmed that under a low T_A_ body temperature was lower in the Low-SA group than High-SA group at 1h after administration of 2, 3 or 4 mg/kg MDMA, but not after 1 or 5 mg/kg. This pattern was similar under a high T_A_, where temperature was lower in the Low-SA group than High-SA group after 2 mg/kg (1h after the session start), 3 mg/kg (1-2h), 4 mg/kg (1-2h) and 5 mg/kg (2h). Thus, a higher level of MDMA self-administration was associated with a blunted hypothermic response across ambient temperatures.

Additionally, for the Low-SA group, post hoc comparisons confirmed that body temperature was lower under a low T_A_ than a hot T_A_ when animals were dosed with 1 mg/kg (Pre, 1 h), 2 mg/kg (1 h), 3 mg/kg (1 h), 4 mg/kg (1-3 h) or 5 mg/kg (1-3 h) of MDMA. However, for the High-SA group, temperature was significantly lower under a low T_A_ than a hot T_A_ only after 3 mg/kg (1-2 h), 4 mg/kg (1-3 h) or 5 mg/kg (1-3 h) of MDMA. Thus, the body temperature responses of the High-SA animals were less impacted by the ambient temperature.

Lastly, body temperature for the Low-SA group was lower than that after the lowest dose (1 mg/kg) under the low T_A_ for most time points (all except 2 mg/kg at Pre and 3 mg/kg at Pre and 3 h), but fewer differences from the 1 mg/kg dose were observed under the high T_A_ (only 4-5 mg/kg at Pre and 2-3 mg/kg at 1 h). Body temperature for the High-SA group was lower than that after the lowest dose under the low T_A_ for fewer time points (2-5 mg/kg at 1 h and 4-5 mg/kg at 2 h as well as 5 mg/kg at 3 h), but under the high T_A_ this was the case only for 4 mg/kg at the pretreatment baseline. Thus, the effect of dose was blunted in the High-SA group as compared to the Low-SA group.

##### 3.1.6.2 Locomotor Activity

The self-admin groups did not exhibit differential activity responses to MDMA (**Figure 7**). The ANOVA confirmed that there were no significant effects of Group or of Group in interaction with time, dose or ambient temperature. However, there were main effects of Dose (F(4.52) = 6.94; p = 0.0001), T_A_ (F_(1.13)_ = 4.88; p < 0.05), and time (F(3.39) = 31.08; p < 0.0001), as well as interactions between Dose and T_A_ (F_(4,52)_ = 4.07; p < 0.01), Dose and time (F_(12,156)_ = 10.10; p < 0.0001), Dose and T_A_ (F_(4,52)_ = 4.07; p < 0.01), time and T_A_ (F_(3,39)_ = 6.01; p < 0.01). There was a trend toward a three way interaction between Dose, T_A_ and time (F_(12,156)_ = 1.69; p = 0.073). Post hoc comparisons (collapsed across self-admin groups) confirmed that activity was higher under a Low T_A_ than a High T_A_ after the three lowest doses both pre-session and in the last 2 hours of the session, that activity increased as a function of dose principally in the 1^st^ hour of the session, and that within-session activity generally decreased over the course of the session for the three lowest doses while at the highest two doses activity showed an inverted-U pattern with a first-hour peak.

### 3.2 Experiment 2: ICSS

The intra-cranial self-stimulation reward threshold was affected by both the ambient temperature (T_A_) and MDMA (**Figure 8**). The ANOVA confirmed significant main effects of T_A_ (F_1,9_=29.10; p<0.0005) and MDMA (F_1,9_=25.70; p<0.001) but not the interaction of factors (F_1,39_=0.39; p=0.55). Post-hoc comparisons confirmed that the threshold was higher than each other condition when vehicle was administered at 30°C and *lower* than every other condition when MDMA was administered at 18°C.

**Figure 8:**
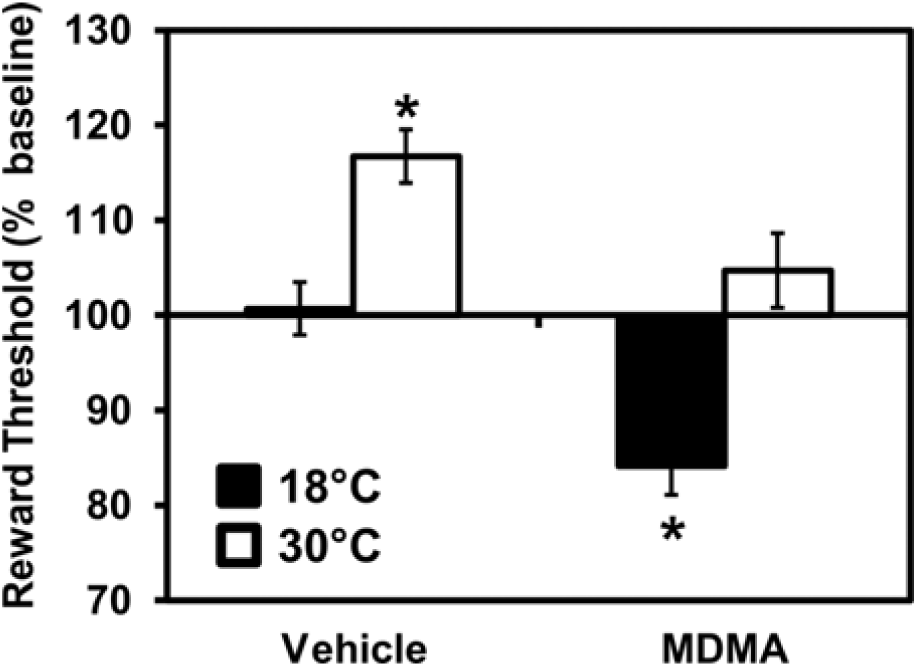
Mean (N=10; ±SEM) ICSS thresholds obtained after treatment with 2.5 mg/kg MDMA, or saline vehicle, s.c., under 18°C or 30°C ambient temperature conditions. Thresholds are expressed as a percentage of individuals’ pre-treatment baseline obtained the same day under 24°C ambient temperature. A significant difference from all other treatment conditions is indicated by *.

## 4. Discussion

### 4.1 High ambient temperature facilitates MDMA IVSA acquisition

The result of this study showed that acquisition of intravenous self-administration (IVSA) of MDMA in rats is affected by the ambient temperature (T_A_) under which animals are trained, with higher temperatures resulting in higher drug intake. The findings are consistent with the prior report of Cornish and colleagues (2003) who found that an *established* MDMA IVSA pattern was increased when the T_A_ was transiently elevated to 30°C. However in this study the increased MDMA IVSA was limited to the initial acquisition and maintenance phases. Raising the room temperature to 30°C after 30 sessions under 20°C *lowered* drug intake, although it did so inconsistently. The difference may be due to the development of tolerance to the intake-enhancing effects of high T_A_ as a consequence of extended low-level MDMA exposure. Alternately it could be proactive interference from the extended period of IVSA training under low T_A_ wherein MDMA as a reinforcer appeared to be less efficacious. Interestingly, the data contrast with another prior investigation which found no difference in the *acquisition* of MDMA self-administration under 23°C versus 32°C ambient temperature [63]. In that study however the responding in the drug groups only differed from the vehicle groups after 15 sessions of acquisition, thus the prior lever training in the present study may be critical. Also, it’s unclear at this time whether or not the small differences in temperatures used in these prior reports (30°C vs 32°C) is substantive. However, with the current study it now appears that under the right conditions, elevated T_A_ can indeed enhance the acquisition of MDMA IVSA. This study therefore provides the first evidence that intoxication with MDMA in a high T_A_ environment, such as at a rave or dance club, may increase the risk for transitioning to a compulsive use pattern.

Although a relatively low number of infusions were self-administered relative to more efficacious stimulant drugs using similar procedures [51, 56], the pattern is consistent with self-administration. Most importantly, drug-associated lever pressing was 83-84% of all responding during acquisition and maintenance, and 87-89% in the temperature swap condition, across both groups. In contrast, our prior study [51] using similar pellet pre-training procedures in a group of rats permitted to self-administer saline under a FR2 contingency found that lever discrimination declined to 40% saline-associated lever pressing after 15 sessions (12 post-food restriction). This differs from the stability observed in the present study up to 49 IVSA sessions. In addition, our prior study found that Sprague-Dawley rats will IVSA about half as many infusions of the entactogen cathinone mephedrone, particularly under high per-infusion training conditions compared with Wistar rats. Accounting for this strain difference the cumulative MDMA dose self-administered in this study is what might be predicted from studies conducted subsequently in Wistar rats [36, 64]. Finally, the increase in responding that was observed during saline substitutions is most parsimoniously interpreted as drug-seeking behavior; a similar pattern for saline relative to a lower training dose was observed in two subsequent studies with male and female Wistar rats subjected to a more extensive dose-substitution procedure, post-acquisition [36, 64].

### 4.2 Within-session hypothermia during MDMA IVSA even under high ambient temperature

These data are the first to fully explicate the MDMA poikilothermic body temperature response [39-41] under intravenous self-administration conditions. A prior study [65] showed that wire caging could block hyperthermia produced by 30 mg/kg s.c. in an acrylic cage. Thus, the predicted effect in operant boxes with metal flooring was for a decrease in body temperature even under 30°C T_A_ since this is still below normal rodent body temperature. The magnitude of the hypothermic response was moderately reduced in the animals acquiring IVSA under hot T_A_ conditions but was still present when this group was swapped to self-administer at the colder temperature (**Figure 4**). The attenuation of the body temperature response developed gradually throughout the acquisition and stable maintenance of MDMA IVSA, thus it appears to be a consequence of cumulative MDMA exposure rather than a condition which itself facilitates increased self-administration. It therefore is unlikely that a change in body temperature has a significant motivational contribution to the establishment of MDMA IVSA, as hypothesized to motivate this study.

The body temperature results differ from a similar, prior study [63] which found no decrease in body temperature associated with MDMA IVSA. Indeed a modest post-session *elevation* in rectal temperature, that was equivalent between MDMA and Vehicle IVSA groups, was observed at each ambient temperature. Since that study involved 2 hr sessions and the temperature sampling was only prior to and after the session, it is possible that much of the dynamic response to self-administered MDMA went undetected (for anticipated time-course see [47, 51] as well as the present data). It is also the case that potential effects of stress associated with rectal sampling of body temperature [66-68] have not been well delineated and may have obscured any drug-related effects in the prior study. Glucocorticoid responses to stress can affect many behaviors in addition to thermoregulation. The lower-ambient group in the study by Feduccia et al. also contrasts with the significant hypothermia reported by the same laboratory in Reveron et al. (2010) and the study of Cornish et al. (2003) did not attempt to measure body temperature. Thus the current data provide a unique window on the thermoregulatory response to MDMA when it is self-administered, i.v..

### 4.3 High ambient temperature lowers reward thresholds

An alternate hypothesis for why high T_A_ caused enhanced IVSA acquisition is that under this condition more drug is needed to lower reward thresholds to the degree necessary for reinforcement. In this study the intracranial self-stimulation reward threshold was increased by high ambient temperature and MDMA was less effective at lowering reward thresholds below the normal-ambient baseline under 30°C versus 18°C T_A_. This suggests that the aversive nature of the higher ambient temperature may have facilitated the establishment of MDMA IVSA in two ways. First, the high ambient temperature increased reward threshold in animals after saline challenge which indicates a negative state which might drive increased drug seeking/taking. Second, the ability of MDMA to lower reward thresholds to a pro-reward state was lessened in the higher T_A_ environment. This suggests that more MDMA may have been required under these conditions before a positive subjective state was attained.

### 4.4 Relatively high levels of MDMA IVSA intake are associated with a blunting of the thermoregulatory response to MDMA, but not to the locomotor activity response

One major interpretive limitation with the present study is that the MDMA dose was self-selected during IVSA. Thus, it is possible that if animals were exposed to high amounts of MDMA in cold T_A_ or low amounts in high T_A_ (e.g., by changing the per-infusion dose) results may differ. Nevertheless, the post-IVSA experiment in which non-contingent doses were administered i.v. demonstrates the dose- and ambient temperature-dependency relationships for the thermoregulatory and locomotor responses reasonably conclusively. Changes in locomotion produced by the 5 mg/kg MDMA, i.v., condition were similar to the magnitude of changes produced by vapor inhalation or i.p. injection of METH, MDPV, or mephedrone [47-49, 69]. The magnitude of temperature reduction after this MDMA dose was also similar to the maximum effects observed for i.p. doses of mephedrone or MDMA [47, 48]. Importantly, these experiments also confirmed a thermoregulatory (but not locomotor) tolerance induced by relatively greater prior exposure to MDMA as a consequence of self-selected doses during IVSA. This underscores the gradual development of thermoregulatory tolerance to MDMA that was observed in the Hot-trained group through the course of the IVSA experiments.

### 4.5 Conclusion

This study confirms that the acquisition of MDMA self-administration is enhanced in higher ambient temperature. Since this was a rodent model featuring intravenous administration rather than the bolus oral consumption typical of human use, extrapolation must be cautious. Nevertheless, the similarity of effect of oral and injected MDMA on thermoregulatory disruption in one animal model [52, 53] suggests route of administration may not be qualitatively different for MDMA. Furthermore, this study emphasizes that the ongoing experience of higher versus lower levels of self-administration have lasting consequences on individual propensity to self-administer under future conditions. No support was provided for the hypothesis that the effects of higher ambient temperature were conveyed by a blunted initial hypothermic response to MDMA; any blunting developed as a consequence of ongoing higher levels of self-administration. The ICSS experiment suggested that the higher intake of MDMA in the hot environment may have been driven by the increased brain reward thresholds caused by the hot environment, and not by any increased rewarding efficacy associated with a given dose of MDMA.

## Authorship and Acknowledgements

This work was originally conceptualized by MAT and the specific experimental designs were determined by MAT and SMA with additional input from PKH. Studies were conducted by SMA and PKH. Data analysis, figure creation and manuscript drafting was by MAT and SMA. All authors critically reviewed content and approved the manuscript for publication. This work was supported by USPHS grants DA024105 and DA042211; the NIH/NIDA had no further role in study design; in the collection, analysis and interpretation of data; in the writing of the report; or in the decision to submit the paper for publication. This is manuscript #28066 from The Scripps Research Institute.

## Literature Cited

1. Peroutka, S.J., Incidence of recreational use of 3,4-methylenedimethoxymethamphetamine (MDMA, “ecstasy”) on an undergraduate campus [letter]. N Engl J Med, 1987. 317(24): p. 1542-3.

2. Pope, H.G., Jr., M. Ionescu-Pioggia, and K.W. Pope, Drug use and life style among college undergraduates: a 30-year longitudinal study. Am J Psychiatry, 2001. 158(9): p. 1519-21.

3. Schuster, P., et al., Is the use of ecstasy and hallucinogens increasing? Results from a community study. Eur Addict Res, 1998. 4(1-2): p. 75-82.

4. Johnston, L.D., et al., Monitoring the Future national survey results on drug use, 1975-2011. Volume I, Secondary school students 2012, University of Michigan: Ann Arbor. p. 760.

5. Johnston, L.D., et al., Monitoring the Future national survey results on drug use, 1975-2011. Volume II, College students and adults ages 19-50. 2012, University of Michigan: Ann Arbor. p. 314.

6. Johnston, L.D., et al. 2012 Overview Key Findings on Adolescent Drug Use. 2013 [cited 2013 03/21/2013]; Available from: http://www.monitoringthefuture.org.

7. Kerbage, H. and S. Richa, Non-Antidepressant Long-Term Treatment in Post-Traumatic Stress Disorder (PTSD). Curr Clin Pharmacol, 2013.

8. Oehen, P., et al., A randomized, controlled pilot study of MDMA (+/-3,4-Methylenedioxymethamphetamine)-assisted psychotherapy for treatment of resistant, chronic Post-Traumatic Stress Disorder (PTSD). J Psychopharmacol, 2013. 27(1): p. 40-52.

9. Cukor, J., et al., Emerging treatments for PTSD. Clin Psychol Rev, 2009. 29(8): p. 715-26.

10. Doblin, R., A clinical plan for MDMA (Ecstasy) in the treatment of posttraumatic stress disorder (PTSD): partnering with the FDA. J Psychoactive Drugs, 2002. 34(2): p. 185-94.

11. Mithoefer, M.C., et al., Durability of improvement in post-traumatic stress disorder symptoms and absence of harmful effects or drug dependency after 3,4-methylenedioxymethamphetamine-assisted psychotherapy: a prospective long-term follow-up study. J Psychopharmacol, 2013. 27(1): p. 28-39.

12. Mithoefer, M.C., et al., The safety and efficacy of {+/-}3,4-methylenedioxymethamphetamine-assisted psychotherapy in subjects with chronic, treatment-resistant posttraumatic stress disorder: the first randomized controlled pilot study. J Psychopharmacol, 2011. 25(4): p. 439-52.

13. Johnston, L.D., et al., Monitoring the Future national survey results on drug use, 1975-2005. Volume II, College students and adults ages 19-40, in (NIH Publication No. 06-5884). 2006, National Institute on Drug Abuse: Bethesda, MD.

14. Thomasius, R., et al., Mental disorders in current and former heavy ecstasy (MDMA) users. Addiction, 2005. 100(9): p. 1310-9.

15. Cottler, L.B., et al., Ecstasy abuse and dependence among adolescents and young adults: applicability and reliability of DSM-IV criteria. Hum Psychopharmacol, 2001. 16(8): p. 599-606.

16. de Almeida, S.P., M. Garcia-Mijares, and M.T. Silva, Patterns of ecstasy use and associated harm: results of a Brazilian online survey. Subst Use Misuse, 2009. 44(14): p. 2014-27.

17. Hurault de Ligny, B., et al., Early loss of two renal grafts obtained from the same donor: role of ecstasy? Transplantation, 2005. 80(1): p. 153-6.

18. Jansen, K.L., Ecstasy (MDMA) dependence. Drug Alcohol Depend, 1999. 53(2): p. 121-4.

19. Kouimtsidis, C., et al., Neurological and psychopathological sequelae associated with a lifetime intake of 40,000 ecstasy tablets. Psychosomatics, 2006. 47(1): p. 86-7.

20. Beardsley, P.M., R.L. Balster, and L.S. Harris, Self-administration of methylenedioxymethamphetamine (MDMA) by rhesus monkeys. Drug Alcohol Depend, 1986. 18(2): p. 149-57.

21. Lamb, R.J. and R.R. Griffiths, Self-injection of d,1-3,4-methylenedioxymethamphetamine (MDMA) in the baboon. Psychopharmacology (Berl), 1987. 91(3): p. 268-72.

22. Fantegrossi, W.E., et al., 3,4-Methylenedioxymethamphetamine (MDMA, “ecstasy”) and its stereoisomers as reinforcers in rhesus monkeys: serotonergic involvement. Psychopharmacology (Berl), 2002. 161(4): p. 356-64.

23. Fantegrossi, W.E., et al., Behavioral and neurochemical consequences of long-term intravenous self-administration of MDMA and its enantiomers by rhesus monkeys. Neuropsychopharmacology, 2004. 29(7): p. 1270-81.

24. Lile, J.A., J.T. Ross, and M.A. Nader, A comparison of the reinforcing efficacy of 3,4-methylenedioxymethamphetamine (MDMA, “ecstasy”) with cocaine in rhesus monkeys. Drug Alcohol Depend, 2005. 78(2): p. 135-40.

25. De La Garza, R., 2nd, K.R. Fabrizio, and A. Gupta, Relevance of rodent models of intravenous MDMA self-administration to human MDMA consumption patterns. Psychopharmacology (Berl), 2006: p. 1-10.

26. Ratzenboeck, E., et al., Reinforcing effects of MDMA (“ecstasy”) in drug-naive and cocaine-trained rats. Pharmacology, 2001. 62(3): p. 138-44.

27. Reveron, M.E., E.Y. Maier, and C.L. Duvauchelle, Behavioral, thermal and neurochemical effects of acute and chronic 3,4-methylenedioxymethamphetamine (“Ecstasy”) self-administration. Behavioural Brain Research, 2010. 207(2): p. 500-507.

28. Daniela, E., et al., Effect of SCH 23390 on (+/-)-3,4-methylenedioxymethamphetamine hyperactivity and self-administration in rats. Pharmacol Biochem Behav, 2004. 77(4): p. 745-50.

29. Daniela, E., D. Gittings, and S. Schenk, Conditioning Following Repeated Exposure to MDMA in Rats: Role in the Maintenance of MDMA Self-Administration. Behav Neurosci, 2006. 120(5): p. 1144-50.

30. Schenk, S., et al., Development, maintenance and temporal pattern of self-administration maintained by ecstasy (MDMA) in rats. Psychopharmacology (Berl), 2003. 169(1): p. 21-7.

31. Schenk, S., et al., MDMA self-administration in rats: acquisition, progressive ratio responding and serotonin transporter binding. Eur J Neurosci, 2007. 26(11): p. 3229-36.

32. Bird, J. and S. Schenk, Contribution of impulsivity and novelty-seeking to the acquisition and maintenance of MDMA self-administration. Addict Biol, 2012.

33. Colussi-Mas, J., et al., Drug seeking in response to a priming injection of MDMA in rats: relationship to initial sensitivity to self-administered MDMA and dorsal striatal dopamine. Int J Neuropsychopharmacol, 2010. 13(10): p. 1315-27.

34. Dalley, J.W., et al., Enduring deficits in sustained visual attention during withdrawal of intravenous methylenedioxymethamphetamine self-administration in rats: results from a comparative study with d-amphetamine and methamphetamine. Neuropsychopharmacology, 2007. 32(5): p. 1195-206.

35. Reveron, M.E., E.Y. Maier, and C.L. Duvauchelle, Experience-dependent changes in temperature and behavioral activity induced by MDMA. Physiol Behav, 2006.

36. Vandewater, S.A., K.M. Creehan, and M.A. Taffe, Intravenous self-administration of entactogen-class stimulants in male rats. Neuropharmacology, 2015. 99: p. 538-545.

37. Cornish, J.L., et al., Heat increases 3,4-methylenedioxymethamphetamine self-administration and social effects in rats. Eur J Pharmacol, 2003. 482(1-3): p. 339-41.

38. Banks, M.L., et al., Effects of ambient temperature on the relative reinforcing strength of MDMA using a choice procedure in monkeys. Psychopharmacology (Berl), 2008. 196(1): p. 63-70.

39. Malberg, J.E. and L.S. Seiden, Small changes in ambient temperature cause large changes in 3,4-methylenedioxymethamphetamine (MDMA)-induced serotonin neurotoxicity and core body temperature in the rat. J Neurosci, 1998. 18(13): p. 5086-94.

40. Dafters, R.I., Effect of ambient temperature on hyperthermia and hyperkinesis induced by 3,4-methylenedioxymethamphetamine (MDMA or “ecstasy”) in rats. Psychopharmacology (Berl), 1994. 114(3): p. 505-8.

41. Dafters, R.I., Hyperthermia following MDMA administration in rats: effects of ambient temperature, water consumption, and chronic dosing. Physiol Behav, 1995. 58(5): p. 877-82.

42. Farrell, W.J. and J.R. Alberts, Rat behavioral thermoregulation integrates with nonshivering thermogenesis during postnatal development. Behav Neurosci, 2007. 121(6): p. 1333-41.

43. Dymond, K.E. and J.E. Fewell, Coordination of autonomic and behavioral thermoregulatory responses during exposure to a novel stimulus in rats. Am J Physiol, 1998. 275(3 Pt 2): p. R673-6.

44. Briese, E., Circadian body temperature rhythm and behavior of rats in thermoclines. Physiol Behav, 1986. 37(6): p. 839-47.

45. Gordon, C.J., Thermal biology of the laboratory rat. Physiol Behav, 1990. 47(5): p. 963-91.

46. Gordon, C.J., et al., Dynamics of behavioral thermoregulation in the rat. Am J Physiol, 1991. 261(3 Pt 2): p. R705-11.

47. Miller, M.L., et al., Changes in ambient temperature differentially alter the thermoregulatory, cardiac and locomotor stimulant effects of 4-methylmethcathinone (mephedrone). Drug Alcohol Depend, 2013. 127(1-3): p. 248-53.

48. Wright, M.J., Jr., et al., Effect of ambient temperature on the thermoregulatory and locomotor stimulant effects of 4-methylmethcathinone in Wistar and Sprague-Dawley rats. PLoS One, 2012. 7(8): p. e44652.

49. Aarde, S.M., et al., In vivo potency and efficacy of the novel cathinone alpha-pyrrolidinopentiophenone and 3,4-methylenedioxypyrovalerone: self-administration and locomotor stimulation in male rats. Psychopharmacology (Berl), 2015. 232(16): p. 3045-55.

50. Nguyen, J.D., et al., Inhaled delivery of Delta-tetrahydrocannabinol (THC) to rats by e-cigarette vapor technology. Neuropharmacology, 2016. 109: p. 112-120.

51. Aarde, S.M., et al., Mephedrone (4-methylmethcathinone) supports intravenous self-administration in Sprague-Dawley and Wistar rats. Addict Biol, 2013. 18(5): p. 786-99.

52. Crean, R.D., S.A. Davis, and M.A. Taffe, Oral administration of (+/-)3,4-methylenedioxymethamphetamine and (+)methamphetamine alters temperature and activity in rhesus macaques. Pharmacol Biochem Behav, 2007.

53. Crean, R.D., et al., Effects of (+/-)3,4-methylenedioxymethamphetamine, (+/-)3,4-methylenedioxyamphetamine and methamphetamine on temperature and activity in rhesus macaques. Neuroscience, 2006. 142(2): p. 515-25.

54. Taffe, M.A., Delta9-Tetrahydrocannabinol attenuates MDMA-induced hyperthermia in rhesus monkeys. Neuroscience, 2012. 201: p. 125-33.

55. Taffe, M.A., et al., Hyperthermia induced by 3,4-methylenedioxymethamphetamine in unrestrained rhesus monkeys. Drug Alcohol Depend, 2006. 82(3): p. 276-81.

56. Aarde, S.M., et al., The novel recreational drug 3,4-methylenedioxypyrovalerone (MDPV) is a potent psychomotor stimulant: self-administration and locomotor activity in rats. Neuropharmacology, 2013. 71: p. 130-40.

57. Aarde, S.M., et al., Binge-like acquisition of 3,4-methylenedioxypyrovalerone (MDPV) self-administration and wheel activity in rats. Psychopharmacology (Berl), 2015. 232(11): p. 1867-77.

58. Miller, M.L., et al., Reciprocal inhibitory effects of intravenous d-methamphetamine self-administration and wheel activity in rats. Drug Alcohol Depend, 2012. 121(1-2): p. 90-6.

59. Schenk, S., et al., Profile of MDMA Self-Administration from a Large Cohort of Rats: MDMA Develops a Profile of Dependence with Extended Testing. Journal of Drug and Alcohol Research, 2012. 1(1): p. 1-6.

60. Macenski, M.J. and R.A. Meisch, Cocaine self-administration under conditions of restricted and unrestricted food access. Exp Clin Psychopharmacol, 1999. 7(4): p. 324-37.

61. Certel, K., et al., Restricted patterning of vestigial expression in Drosophila wing imaginal discs requires synergistic activation by both Mad and the drifter POU domain transcription factor. Development, 2000. 127(14): p. 3173-83.

62. Collins, G.T., et al., Effects of dopamine D(2)-like receptor agonists in mice trained to discriminate cocaine from saline: influence of feeding condition. Eur J Pharmacol, 2014. 729: p. 123-31.

63. Feduccia, A.A., N. Kongovi, and C.L. Duvauchelle, Heat increases MDMA-enhanced NAcc 5-HT and body temperature, but not MDMA self-administration. Eur Neuropsychopharmacol, 2010. 20(12): p. 884-94.

64. Creehan, K.M., S.A. Vandewater, and M.A. Taffe, Intravenous self-administration of mephedrone, methylone and MDMA in female rats. Neuropharmacology, 2015. 92: p. 90-7.

65. Gordon, C.J. and L. Fogelson, Metabolic and thermoregulatory responses of the rat maintained in acrylic or wire-screen cages: implications for pharmacological studies. Physiol Behav, 1994. 56(1): p. 73-9.

66. McGivern, R.F., D.G. Zuloaga, and R.J. Handa, Sex differences in stress-induced hyperthermia in rats: restraint versus confinement. Physiol Behav, 2009. 98(4): p. 416-20.

67. Aydin, C., C.E. Grace, and C.J. Gordon, Effect of physical restraint on the limits of thermoregulation in telemetered rats. Exp Physiol, 2011. 96(11): p. 1218-27.

68. Vidal, C., C. Suaudeau, and J. Jacob, Regulation of body temperature and nociception induced by non-noxious stress in rat. Brain Res, 1984. 297(1): p. 1-10.

69. Nguyen, J.D., et al., Locomotor Stimulant and Rewarding Effects of Inhaling Methamphetamine, MDPV, and Mephedrone via Electronic Cigarette-Type Technology. Neuropsychopharmacology, 2016. 41(11): p. 2759&71.

